# Anticancer Drug Impact on DNA – A Study by Neutron Spectrocopy, Synchrotron-based FTIR and EXAFS

**DOI:** 10.1101/398826

**Authors:** Ana L.M. Batista de Carvalho, Adriana P. Mamede, Asha Dopplapudi, Victoria Garcia Sakai, James Doherty, Mark Frogley, Gianfelice Cinque, Peter Gardner, Diego Gianolio, Luís A.E. Batista de Carvalho, Maria P.M. Marques

## Abstract

A complementary structural and dynamical information on drug-DNA interplay has been achieved at a molecular level, for Pt/Pd-drugs, allowing a better understanding of their pharmacodynamic profile. The interaction of two cisplatin-like dinuclear Pt(II) and Pd(II) complexes with DNA was studied through a multidisciplinary experimental approach, using quasi-elastic neutron scattering (QENS) techniques coupled to synchrotron-based extended X-ray absorption fine structure (SR-EXAFS) and Fourier-Transform Infrared Spectroscopy-Attenuated Total Reflectance (SR-FTIR-ATR). The drug impact on DNA’s dynamical profile, *via* its hydration layer, was provided by QENS, a drug-triggered enhanced mobility having been revealed. Additionally, an onset of anharmonicity was detected for dehydrated DNA, at room temperature. Far- and mid-infrared measurements allowed the first simultaneous detection of the drugs and its primary pharmacological target, as well as the drug-prompted changes in DNA’s conformation that mediate cytotoxicity in DNA extracted from drug-exposed human triple negative breast cancer cells (MDA-MB-231), a low prognosis type of cancer. The local environment of the absorbing Pd(II) and Pt(II) centers in the drugs’ adducts with adenine, guanine and glutathione was attained by EXAFS.

## INTRODUCTION

Cancer is a leading cause of death worldwide, with a growing incidence: 12.7 million people *per* year are diagnosed with cancer, from which more than 50% die from the disease. Early diagnosis and chemotherapeutic treatment can lower this mortality rate by 30 to 40 %, and may lead to the cure of *ca.* 30 % of the patients. Anticancer platinum drugs have been introduced in the clinics since the serendipitous discovery of cisplatin (*cis*-dichlorodiamine platinum(II), c*is*-(NH_3_)_2_PtCl_2_) in the 1970s, which was the first inorganic compound displaying a high antineoplastic activity towards human tumours.(Rosenberg, Vancamp et al., 1965, Rosenberg, VanCamp et al., 1969) The three currently approved platinum agents – cisplatin, carboplatin (*cis*-diamine(1,1-cyclobutanedicarboxylato) platinum(II)) and oxaliplatin ([(1R,2R)-cyclohexane-1,2-diamine](ethanedioato-O,O’) platinum(II)) – are applied worldwide in chemotherapeutic regimes. However, their clinical application is still restricted by dose-limiting deleterious side effects and acquired resistance upon prolonged administration,(Florea & Busselberg, 2011) as well as by lack of specificity against several cancer types (*e.g.* metastatic). Hence, considerable effort has been put into the development of novel metal-based drugs, including cisplatin-like agents, aiming at an improved antitumor efficiency.(Kelland, 2007, Marques, 2013, Wheate, Walker et al., 2010) Among these, polynuclear multifunctional Pt(II) and Pd(II) chelates with flexible biogenic polyamines as bridging ligands have been synthetized by the authors and evaluated as to their effect on several types of human neoplasias, namely the low prognosis, highly metastatic triple negative breast cancer against which a Pd(II) agent (Pd_2_Spm, Spm=spermine, H_2_N(CH_2_)_3_NH(CH_2_)_4_NH(CH_2_)_3_NH_2_) has yielded particularly promising results.(Fiuza, Holy et al., 2011)

A number of studies have been conducted in the last few years on these Pt- and Pd-based complexes, reporting conformational profiles(Batista de Carvalho, Parker et al., 2018, Batista de Carvalho, Marques et al., 2011, Fiuza, Amado et al., 2015, Marques, Valero et al., 2013) and pharmacokinetic/pharmacodynamic behaviour,(Corduneanu, Chiorcea-Paquim et al., 2010, Marques, Gianolio et al., 2015) and the effect on different human cancers(Batista de Carvalho, Medeiros et al., 2016a, Fiuza, Amado et al., 2006, Fiuza et al., 2011, Marques, Girao et al., 2002, Silva, Andersson et al., 2013, Silva, Fiuza et al., 2014, Soares, Fiuza et al., 2007, Tassoni, Bagni et al., 2010, Teixeira, Seabra et al., 2004, Tummala, Diegelman et al., 2010) including their impact on cellular biochemical profile.(Batista de Carvalho, Pilling et al., 2016b, Lamego, Marques et al., 2017, Marques, Batista de Carvalho et al., 2017) Unconventional pathways of cytotoxicity were disclosed, which are recognized to lead to an improved therapeutic efficiency coupled to a lower toxicity. In addition, the data gathered so far suggests distinct mechanisms of action for the Pt(II) *versus* the Pd(II) agents, as well as for the dinuclear polyamine chelates relative to the mononuclear lead drug cisplatin. The activity of this kind of agents was found to be mediated by selective covalent binding of the metal centers to DNA bases (mainly the purines at their most nucleophilic nitrogen atom, N7), yielding long-range intra- and interstrand adducts responsible for cell growth arrest and apoptotic death. Adducts with DNA are formed upon drug activation, according to the following steps: transport into the cell, hydrolysis (in the cytoplasm) by sequential loss of the chloride ligands and aquation, uptake into the nucleus and reaction of the unstable diaquo species with DNA (the drug’s main pharmacological target).(Reedijk, 2003) However, apart from this interplay with DNA, during the pharmacokinetic stage drug interaction may also occur with amine and sulfur groups from proteins and other cellular constituents. In particular, endogenous thiols such as glutathione (L-γ-glutamyl-cysteinyl-glycine (Glu-Cys-Gly), GSH, a key cellular antioxidant) are known to intercept this type of agents owing to the high affinity of the soft Pt(II) and Pd(II) cations towards S-donor ligands,(Chen & Kuo, 2010, Kasherman, Sturup et al., 2009) which may significantly lower the drug’s bioavailability at the main target and lead to acquired resistance. Since the mechanisms underlying drug resistance as well as the molecular basis of cytotoxicity are still not fully understood for these DNA-damaging agents, the present study aims at contributing to their elucidation by monitoring the interaction of spermine Pt(II) and Pd(II) complexes (Pt_2_Spm and Pd_2_Spm) with DNA.

DNA extracted from human triple negative breast cancer cells (MDA-MB-231 cell line) and commercial DNA were used as target models, while cisplatin was taken as a drug reference. Drug concentrations and incubation times were chosen according to previous results, corresponding to an optimal cytotoxic effect towards this particular neoplastic cell line^7^: 4 and 8 μM (up to *ca.* 2xIC_50_ level), for a 48 h exposure time. A multidisciplinary experimental approach was followed, through the application of: (i) Quasi-elastic neutron scattering (QENS), suitable for directly accessing different spatially resolved dynamical processes (at subnanometer lengthscale and subnanosecond time-scale), under distinct conditions, mainly for hydrogen-rich systems;(Foglia, Hazael et al., 2016, Frolich, Gabel et al., 2009, Fujiwara, Araki et al., 2016, Marques et al., 2017) (ii) synchrotron-radiation Fourier-transform Infrared Spectros-copy-Attenuated Total Reflectance (SR-FTIR-ATR), recognized as a cutting-edge non-destructive tool for obtaining spectral signatures of molecular components in biological samples, so as to relate structural to functional data;(Baker, Trevisan et al., 2014, Holman, Bechtel et al., 2010) (iii) synchrotron-based Extended X-ray Absorption Fine Structure (EXAFS) and X-ray Absorption Near-Edge Structure (XANES), that are methods of choice for obtaining detailed information on the local structure of bioinorganic non-crystalline materials.(Obata, Harada et al., 2009, Provost, Bouvet-Muller et al., 2009)

Water supports vital biochemical processes in living organisms, and is responsible for the maintenance of the functional three-dimensional architecture of biopolymers through a tight interplay within their hydration shells.(Laage, Elsaesser et al., 2017) Actually, mobility changes in these hydration layers may affect the biopolymer’s conformational and dynamical profile, which rule biofunctionality. It is a well-recognized fact that water within hydration sheets is retarded with respect to the bulk, although its dynamical properties are not yet completely understood. Hence, elucidation of water dynamics in biological systems and its impact on activity and function is of the utmost relevance in drug development, for an improved understanding of a drugs’ mode of action *via* interaction with all its possible pharmacological targets that may include the water molecules wrapping their conventional receptors. Neutron techniques such as high resolution quasielastic scattering are particularly suited for selectively probing water dynamical behavior, namely within biomolecular hydration shells, at a nano- to picosecond timescale (*ca.* 10^-9^ – 10^-13^ s) and a 1 to 30 Å lengthscale (corresponding to inter- and intramolecular distances, *e.g.* H-bonding) (Mamontov & Chu, 2012), yielding results not achievable by any other methods. QENS measurements at different temperatures allow to characterize the translational and rotational modes of water hydrogens within a biological matrix (*e.g.* biomolecules, cells or even tissues), and determine how this dynamical behavior may be disturbed by the presence of an external entity such as a drug. A few QENS experiments on water dynamics in living cells have been reported,(Frolich et al., 2009) but drug effects were first investigated by the authors in cisplatin-exposed human cancer cells using QENS and inelastic neutron scattering spectroscopy (INS).(Marques et al., 2017)

FTIR spectroscopy is an extremely powerful, sensitive and non-invasive analytical tool for interrogating the chemical composition of biological systems, delivering unique spectral signatures for a particular biomolecule at specific conditions. The synchrotron beam available at the MIRIAM (Multimode InfraRed Imaging And Microspectroscopy) beamline from the Diamond Light Source (DLS, United Kingdom), currently used, provides stable, broadband and extremely bright IR radiation and delivers FTIR data with an unmatched signal-to-noise ratio. In addition, MIRIAM spans the largest infrared spectral range – extending from the near-up to the far-IR (or THz) region – and is one to two orders of magnitude brighter in the mid-/far-IR than any other conventional thermal IR source. Furthermore, the FTIR attenuated total reflection (ATR) mode used in the present measurements allows to directly probe DNA without any particular sample preparation, with the added advantages of high spatial resolution and absence of resonant Mie scattering effects on the collected spectral data.(Chan & Kazarian, 2016) In particular, probing the far-infrared region (< 300 cm^-1^) that comprises the internal vibrational modes of the molecule, enabled to detect H-bonds and weak non-bonding interactions within the nucleic acid, with very high specificity, thus unveiling specific drug-induced conformational changes that underlie cytotoxicity.

EXAFS, in turn, allows the direct observation of metal coordination and elucidation of the local environment of the absorbing metal center in inorganic compounds, particularly when good quality crystals are unavailable, and has been successfully applied to Pt(II) compounds specifically for assessing their degradation in the presence of S-containing molecules.(Obata et al., 2009, Provost et al., 2009) In the present study, synchrotron-based EXAFS/XANES measurements yielded detailed information on the first coordination shell of Pd(II) and Pt(II) within the drugs’ adducts with adenine (A), guanine (G) and glutathione (GSH).

Complementary structural and dynamical information was therefore achieved for the drug-DNA systems presently studied. These results provide a more comprehensive understanding of the pharmacodynamic profile of the dinuclear Pt- and Pd-anticancer agents, at a molecular level, particularly regarding their effect on vital biomolecules through an impact on their hydration water, apart from a direct perturbation of their native conformation. This knowledge is paramount for a rational design of novel metal-based anticancer compounds with an enhanced chemotherapeutic efficiency coupled to lower deleterious side effects.

## RESULTS AND DISCUSSION

The present study aimed at obtaining detailed information on the pharmacokinetic and pharmacodynamic profiles of two polynuclear cisplatin-like Pt(II) and Pd(II) anticancer agents: their impact on DNA’s dynamical and conformational preferences was evaluated by QENS techniques and synchrotron-radiation FTIR-ATR, while their interaction with DNA purine bases, as well as competition from glutathione, were tackled by synchrotron-based EXAFS/XANES measurements (Scheme 1).

**Scheme 1.**
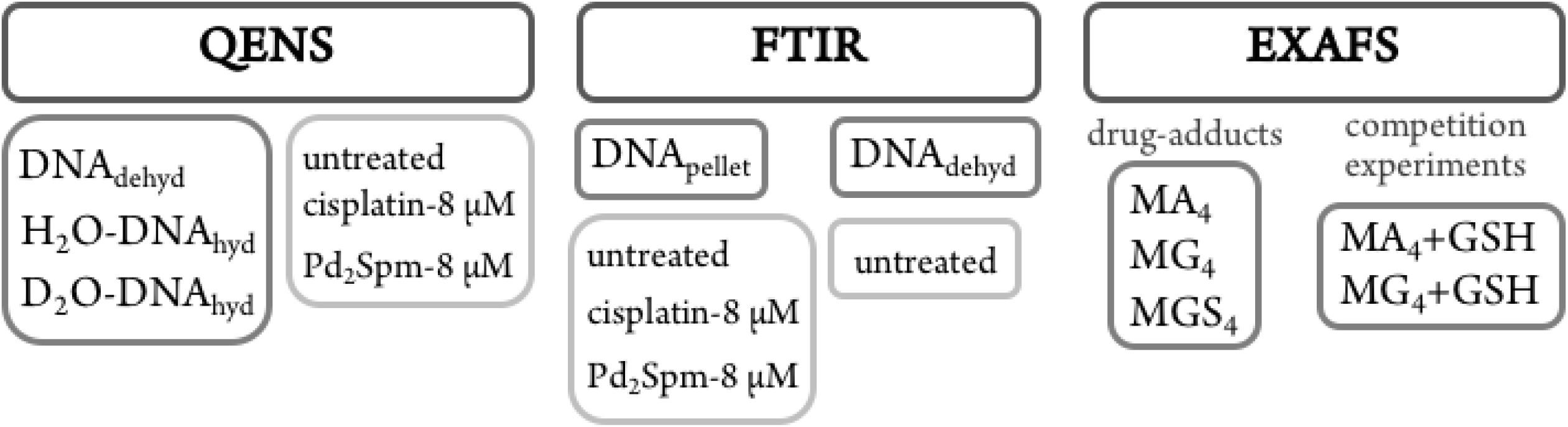
Schematic representation of the samples presently studied and of the techniques used for their analysis. DNA samples: hyd – hydrated; dehyd – dehydrated; pellet – isolated from MDA-MB-231 cells; MA_4_, MG_4_ and MGS_4_ – drug adducts with adenine (A), guanine (G) or glutathione (GSH) (M=Pt or Pd).

### QENS

It is a well-recognized fact that biomolecules are subject to a strict relationship between structure/conformation and activity, as well as between dynamical behavior and function. Hydration is critical for bioactivity, the first hydration shell being an essential part of a biomolecule’s structure, strictly regulating its conformational and dynamical preferences and consequently its physiological role.(Sokolov, Roh et al., 2008, Stadler, Demmel et al., 2016) Regarding DNA, interactions with water are crucial to ensure the native B-conformation of the double helix, thus modulating functionality.(Duboue-Dijon, Fogarty et al., 2016) The dynamics of water within the hydration layer differ from that of bulk water, and are dependent on the biomolecule’s conformational fluctuations as well as on the chemical and topological heterogeneity of its surface (*e.g.* phosphate groups or DNA minor *vs* major grooves). Conversely, even minor variations in hydration water dynamics may prompt significant rearrangements in DNA that can severely distress the normal cellular function and lead to growth inhibition and cell death. An external entity such as a drug may affect the H-bonding network of water in these first hydration shells, both amid water molecules and between water and the biomolecule. Hydration water (as well as intracellular water, in a cellular or tissue matrix) may thus constitute a potential secondary pharmacological target, namely for anticancer chemotherapeutic compounds.(Marques et al., 2017) This concept was currently applied for the elucidation of the drug-target interaction of metal-based agents, which is an innovative approach to monitor pharmacodynamics and better interpret a drug’s mechanism of action, with a view to improve cytotoxicity.

DNA hydration water is mostly adsorbed, in a cooperative way, at the outer double helix surface, with a higher density around the phosphate groups and more ordered near the bases (*ca.* 18-30 water molecules *per* nucleotide(Makarov, Pettitt et al., 2002)). In this first hydration shell, H-bond interactions are stronger than in bulk water and water mobility is significantly reduced, with residence times about 10 times larger.(Beta, Michalarias et al., 2003) Although relaxation within DNA and its hydration layer is slower than for RNA and proteins, a dynamical transition was detected for this nucleic acid at 200-230 K, similarly to other hydrated biopolymers (globular proteins, GFP, RNA). (Chen, Liu et al., 2006, Khodadadi, Roh et al., 2010, Sokolov et al., 2008) Below this temperature, flexibility is significantly reduced and functionality is lost. This transition is thought to be triggered by strong coupling with the hydration water molecules, that undergo mobility changes at the same temperature (*via* H-bonding and translational processes). Consequently, it is strongly affected by the properties of the surrounding medium, which may vary with temperature, pressure, pathological conditions or the presence of non-endogenous compounds such as a drug.(Chen et al., 2006, Wood, 2016) Despite the still poorly understood nature of this dynamical transition, it is generally acknowledged as essential for biological activity.

Building on the success of the first QENS experiment performed by the authors on a drug’s impact in human breast cancer cells, that unveiled a noticeable dose-dependent cisplatin effect on intracellular water dynamics (both for the cytoplasm and hydration layers),(Marques et al., 2017) the present study applied quasi-elastic neutron scattering spectroscopy (with isotope labelling) to probe hydrated DNA, upon chemotherapeutic exposure. This allowed us to detect drug-elicited dynamical changes, that were assessed through variations in the mobility of the labile protons from the macromolecule and its hydration layer – observable within the time- and lengthscales of the OSIRIS spectrometer, *ca.* 4-200 ps and 4-20 Å. DNA samples (prepared from fibrous calf thymus DNA) were measured before and after incubation with either cisplatin or Pd_2_Spm. Both H_2_O- and D_2_O-hydrated samples were probed (DNA_hyd_), with a view to differentiate between the drug effects on: (i) DNA’s exchangeable hydrogens and hydration water; (ii) DNA’s backbone and non-labile hydrogens. The hydration degree was kept constant (r.h. >80%), ensuring that the measured variations in the QENS profiles were solely due to the effect of the drug and not to transitions induced by differences in the macromolecule’s hydration. The H_2_O-DNA_hyd_ samples comprised all dynamical contributions (hydration water, exchangeable and non-exchangeable H’s and DNA’s skeleton), while the slower motions from the non-labile hydrogens and DNA phosphoribose backbone were retrieved from the D_2_O-DNA samples. Dehydrated DNA (DNA_dehyd_) was also analyzed, with and without the tested compounds. Any effect of the experimental protocol on the dynamical behavior of the drug-treated DNA_dehyd_ samples was ruled out by comparing the QENS profiles of DNA_dehyd_ as received (from Sigma) and a corresponding sample prepared in the same way as those exposed to either cisplatin or Pd_2_Spm (see Experimental section). For these dehydrated samples, the main foreseen dynamical processes are the global modes encompassing the macromolecule’s heavy atoms (slow backbone motions) not expected to be detected within the OSIRIS time window. The experiments were performed at 150 and 298 K, that span below and above the dynamical transition temperature reported for B-DNA (222 K(Chen et al., 2006)). In contrast to room temperature, no significant differences were observed between the dynamical profiles of H_2_O- and D_2_O-DNA_hyd_ at 150 K, *i.e.* below the dynamical crossover temperature the hydration layer is more rigid and the H_2_O-hydrated nucleic acid gradually reaches a dynamical profile similar to that of the D_2_O-hydrated molecule (the internal DNA dynamics being undetectable in OSIRIS).

Figure 1 depicts the QENS profiles (Figure 1(A)) and elastic scan plots (elastic intensity *vs* temperature for the whole temperature range probe, Figure 1(B) to (D)) for H_2_O- and D_2_O-DNA_hyd_ as well as for DNA_dehyd_, clearly showing a dynamical transition taking place at *ca.* 225 K for H_2_O-DNA_hyd_ and at *ca.* 260 K for D_2_O-DNA_hyd_. A gentle linear-like temperature dependence was detected below the cross-over temperature, while above it different behaviors were observed for each sample: (i) for D_2_O-DNA_hyd_ a dynamic component was observed, driven by its D_2_O-hydration shell, and an anticipated mobility reduction relative to H_2_O-DNA_hyd_ was verified (Figure 1(B)); (ii) regarding dehydrated DNA (lyophilized DNA, lacking an hydration layer), some dynamical contribution was still detected within the OSIRIS timescale (Figure 1(C) and (D)). This was also evidenced through the QENS profiles for H_2_O-DNA_hyd_ and D_2_O-DNA_hyd_ *versus* DNA_dehyd_ samples (Figure 1(A)). This dynamic behavior currently unveiled for dehydrated DNA is surprising, and not in complete accordance with previous studies that reported an harmonic (temperature independent) profile for dry DNA (commercially available lyophilized sample)(Nickels, Curtis et al., 2012, Sokolov, Grimm et al., 2001) and identified hydration water motions as the sole responsible for the dynamical transition in this system. However, anharmonicity has been formerly reported for dry lysozyme and assigned to methyl group relaxation-like motions.(Roh, 2016, Roh, Novikov et al., 2005). Accordingly, we tentatively ascribe the onset of anharmonicity presently detected for dehydrated DNA to intrinsic relaxation processes, namely rotational motions of CH_3_ as well as of H-bond free NH_2_ groups within the nucleic acid molecule (at the nitrogen bases). In fact, methyls have been recognized as plasticizers of proteins, with a high impact on their dynamics and activity.(Nickels et al., 2012) Although DNA contains a much lower amount of CH_3_ moieties as compared to proteins (one CH_3_ (in thymine) *per* each four nitrogen bases), its effect should be similar even if to a much lower extent. Since methyl groups are symmetric tops (C_3v_ symmetry), they have a very low rotational energy barrier and therefore facilitate the dynamics of biomolecules, namely DNA, even for low hydration levels.

**Figure 1:**
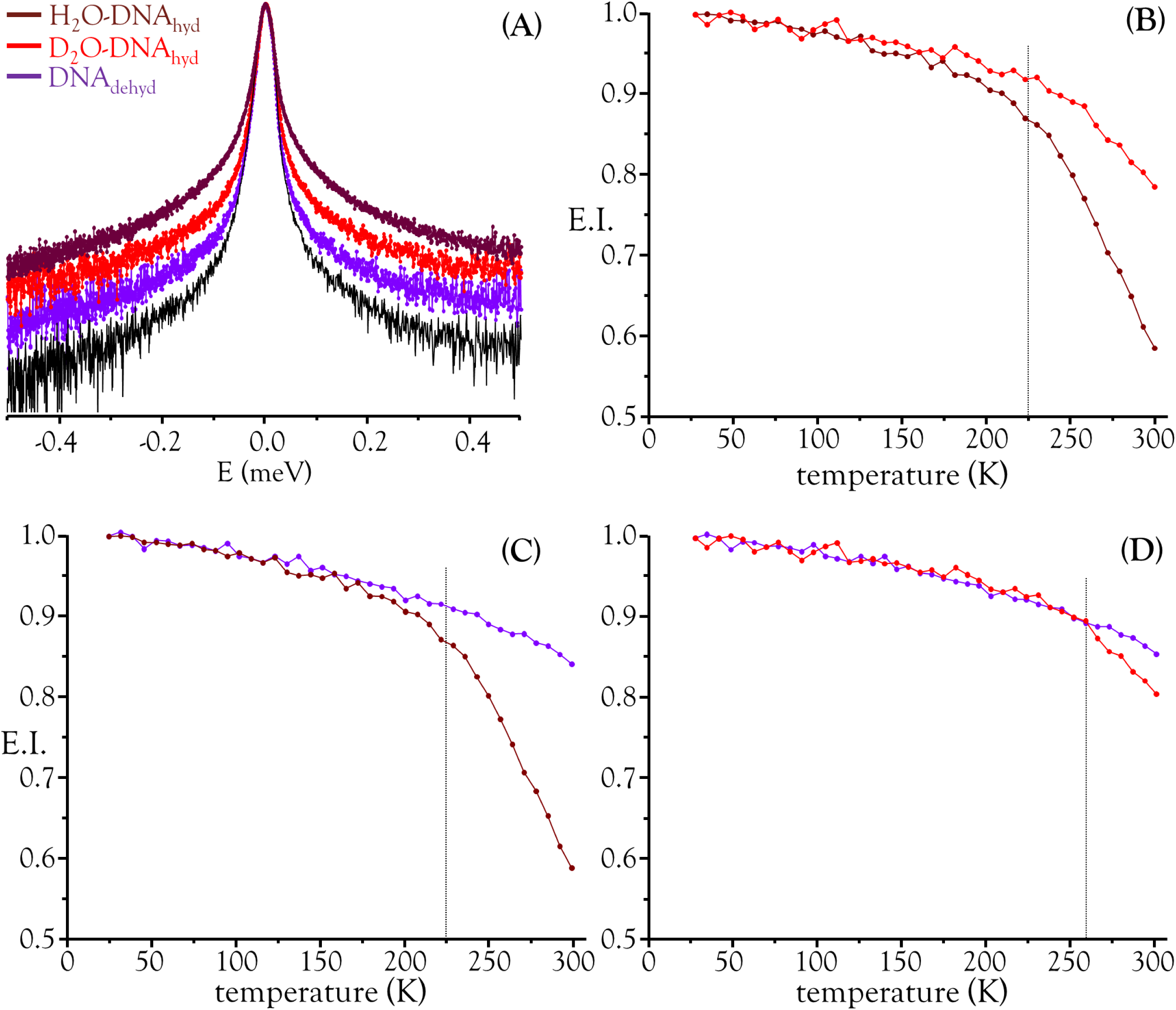
QENS data for hydrated *versus* dehydrated DNA. (A) QENS profiles (for all Q values, at 298 K, logarithmic scale). Elastic scan plots (20-298 K): (B) D_2_O-DNA_hyd_ *vs* H_2_O-DNA_hyd_; (C) DNA_dehyd_ *vs* H_2_O-DNA_hyd_; (D) DNA_dehyd_ *vs* D_2_O-DNA_hyd_. (In (A), the spectra were normalised to maximum peak intensity. The black line represents the instrument resolution).

The drug influence through an effect on DNA’s hydration layer was only detected at 298 K, which appears to indicate that hydration water and DNA motions, in the ps timescale, are not directly coupled at low temperatures. Furthermore, at 150 K (well below the dynamical transition temperature) the hydration shell is considerably less fluid and its dynamics are too slow to be probed with the OSIRIS instrument (Figure 2(A)). Also, no drug effect was found for dehydrated DNA (Figure 2(B)). For the hydrated nucleic acid, drug exposure was found to prompt a faster DNA dynamics, justified by the disruption of the ordered hydration shell and of the native conformation of the nucleic acid upon drug binding, known to affect the native base-pair and base packing arrangements: for drug-incubated DNA, a broader QENS profile was measured (Figure 3(A)), as well as a slight deviation to faster kinetics above the dynamical transition temperature. The effect of cisplatin was found to be somewhat stronger than that of Pd_2_Spm (Figure 3(A) *vs* (B)), which is in accordance with previous studies by vibrational microspectroscopy on these drugs’ influence on the cellular metabolic profile.(Batista de Carvalho et al., 2016b) Interestingly enough, the drug impact on DNA’s hydration layer was not found to affect the biomolecule’s dynamical transition temperature, neither for cisplatin nor for the dinuclear Pd-agent.

**Figure 2:**
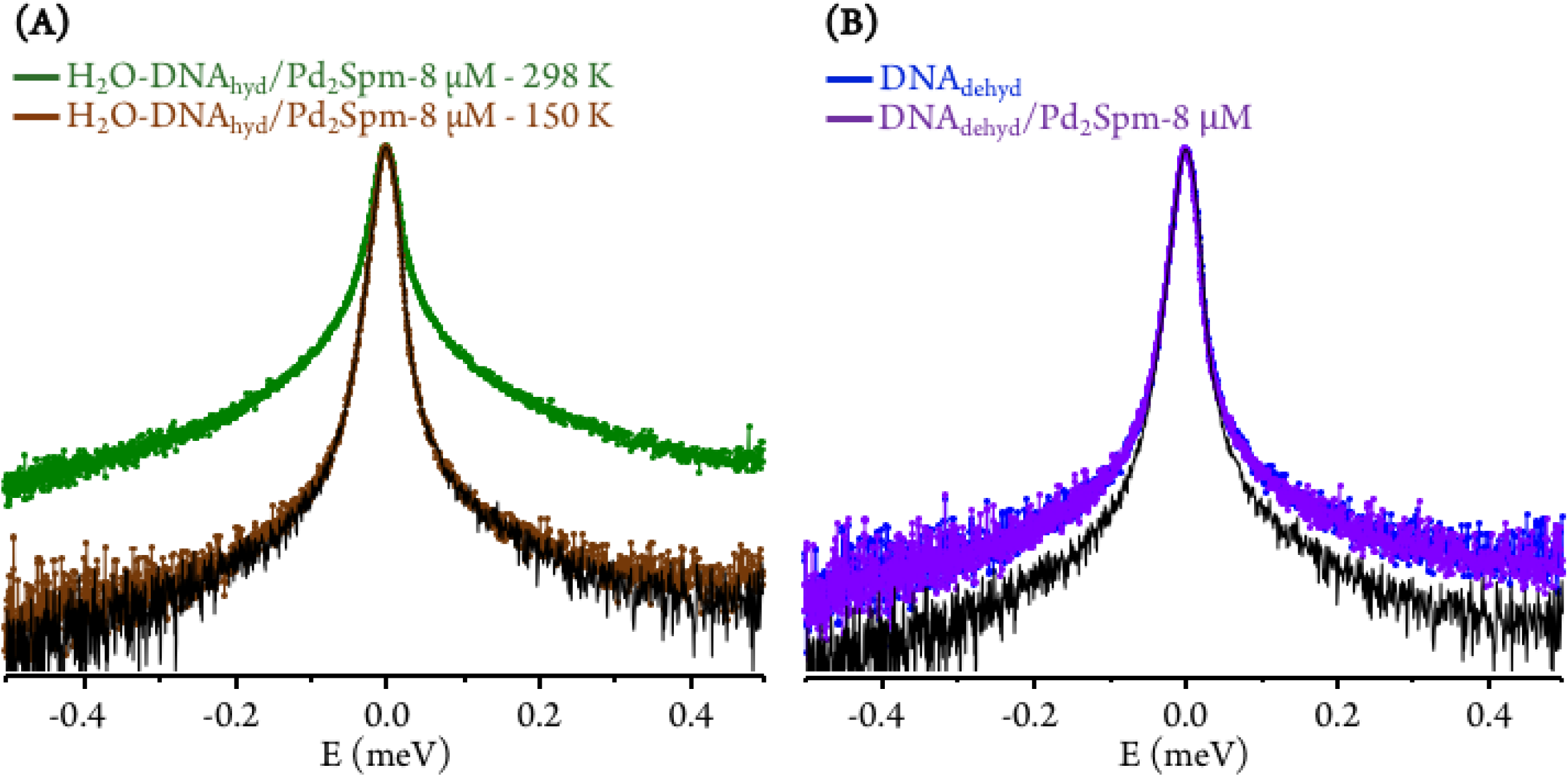
QENS profiles (for all Q values, logarithmic scale): (A) Pd_2_Spm-treated H_2_O-DNA_hyd_, at 150 and 298 K; (B) untreated and Pd_2_Spm-treated DNA_dehyd_, at 298 K. (The spectra were normalized to maximum peak intensity. The black line represents the instrument resolution).

**Figure 3:**
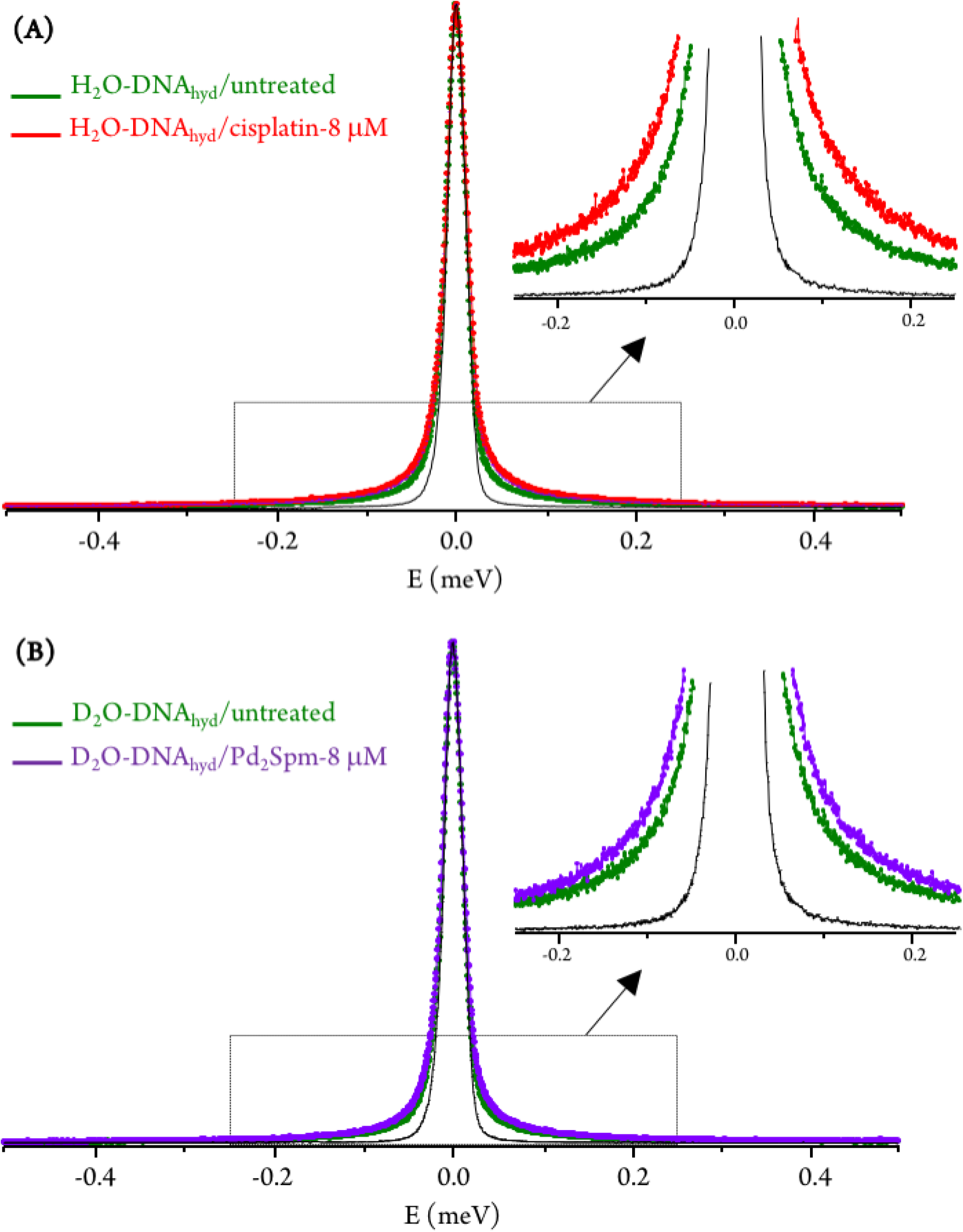
QENS profiles (298 K, for all Q values) for untreated and drug-exposed H_2_O-DNA_hyd_: (A) cisplatin-treated. (B) Pd_2_Spm-treated. (The spectra were normalized to maximum peak intensity. The black line represents the instrument resolution).

No drug effect was found for either D_2_O-hydrated DNA pellet or DNA_dehyd_ (Figure 2(B)), suggesting that the slow motions of the nucleic acid skeleton (backbone dynamics), although expected to be affected by the DNA -binding compounds presently studied, are not detected within the OSIRIS timescale. In addition, deuteration of the hydration shell in D_2_O-DNA_hyd_ samples and the absence of an hydration layer in DNA_dehyd_ appear to hinder a significant drug impact on the macromolecule, evidencing the key role of mobile H’s and hydration water molecules as key mediators in external perturbations to the biomolecule. Nevertheless, these are only tentative conclusions that require confirmation on a higher resolution QENS spectrometer, allowing the detection of the slower dynamical processes (in the nanoscale timescale) that cannot be accessed in OSIRIS.

The neutron diffraction plots obtained for H_2_O-DNA_hyd_ display two noticeable Bragg peaks (at *ca.* 0.35 and 0.55 Å^-1^, detected at both 298 and 150 K, Figure 4) that reflect the ordered structure of the nucleic acid molecule. In the presence of the drug (either cisplatin or Pd_2_Spm), the most intense peak is clearly shifted from 0.35 to 0.3 Å^-1^, which corresponds to a variation in distance of *ca.* 3 Å (from 17.9 to 20.9 Å), that may be related to both the gap between stacked bases (*ca.* 3.3 Å) and the distance between H-bonded base pairs (*ca.* 2.8 to 2.95 Å) in *ds*DNA helices. This effect (found to be independent of temperature) supports a drug-elicited structure disruption of the B-DNA molecule, associated to a perturbation of the nucleic acid’s base packing and pairing arrangements.

**Figure 4:**
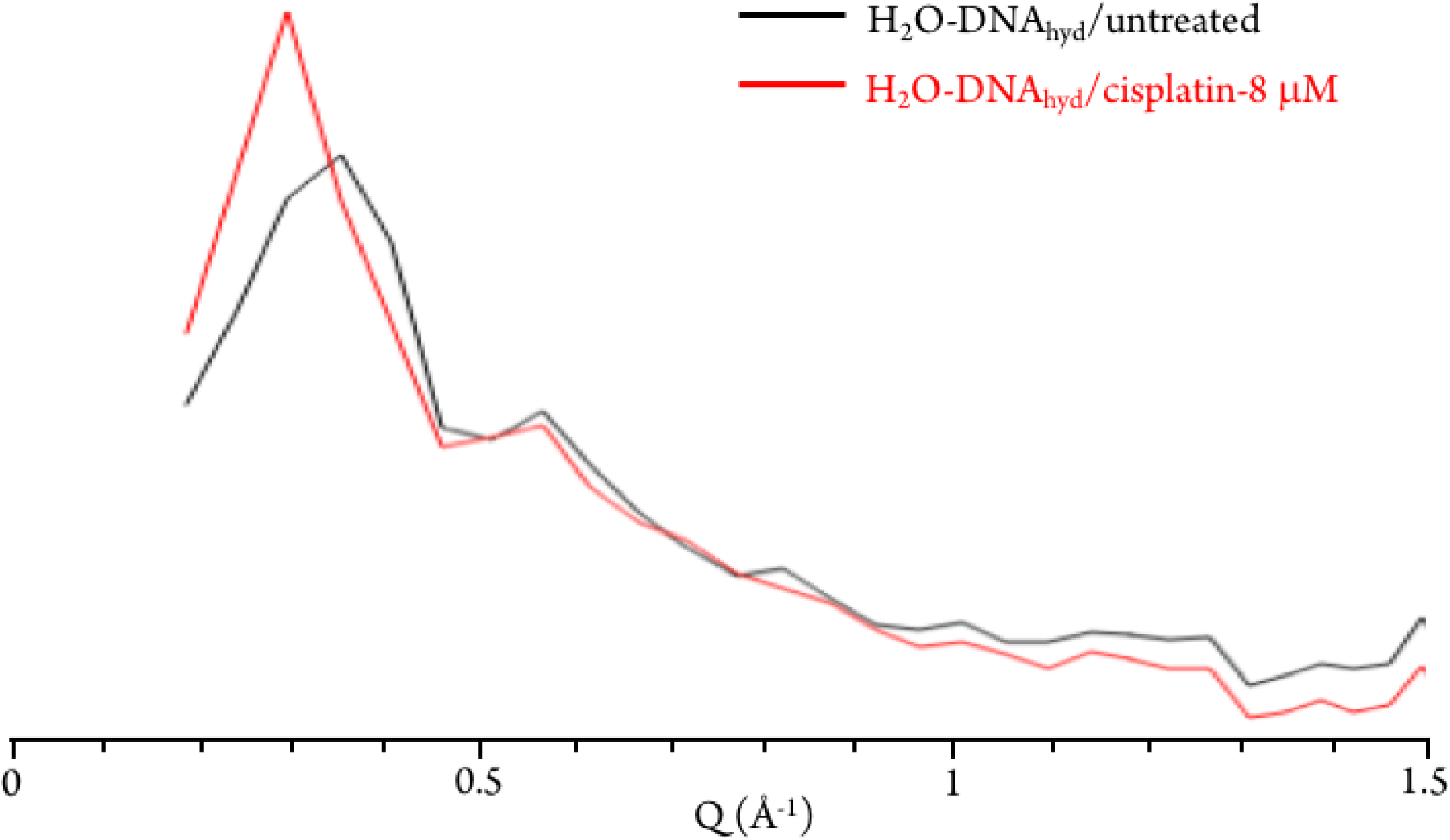
Diffraction plots (at 298 K) for untreated and cisplatin-exposed H_2_O-hydrated DNA.

The experimental data was best fit using one δ-function (elastic component) convoluted with two Lorentzians (quasielastic contributions) (eq. (5), Figure 5). The very slow global motions of the macromolecule are defined by the Delta function (slower than the longest observable time defined by the instrument resolution), while the narrow (Γ_global_) and broader (Γ_local_) Lorentzians represent, respectively: (i) slow motions (Q-dependent) from the biopolymer’s hydration water and H-bond restricted moieties located at the molecule’s surface (side-chain motions); (ii) fast localized motions (Q-independent) ascribed to the rotation of CH_3_ groups and also of amine moieties (not involved in H-bonds).

**Figure 5:**
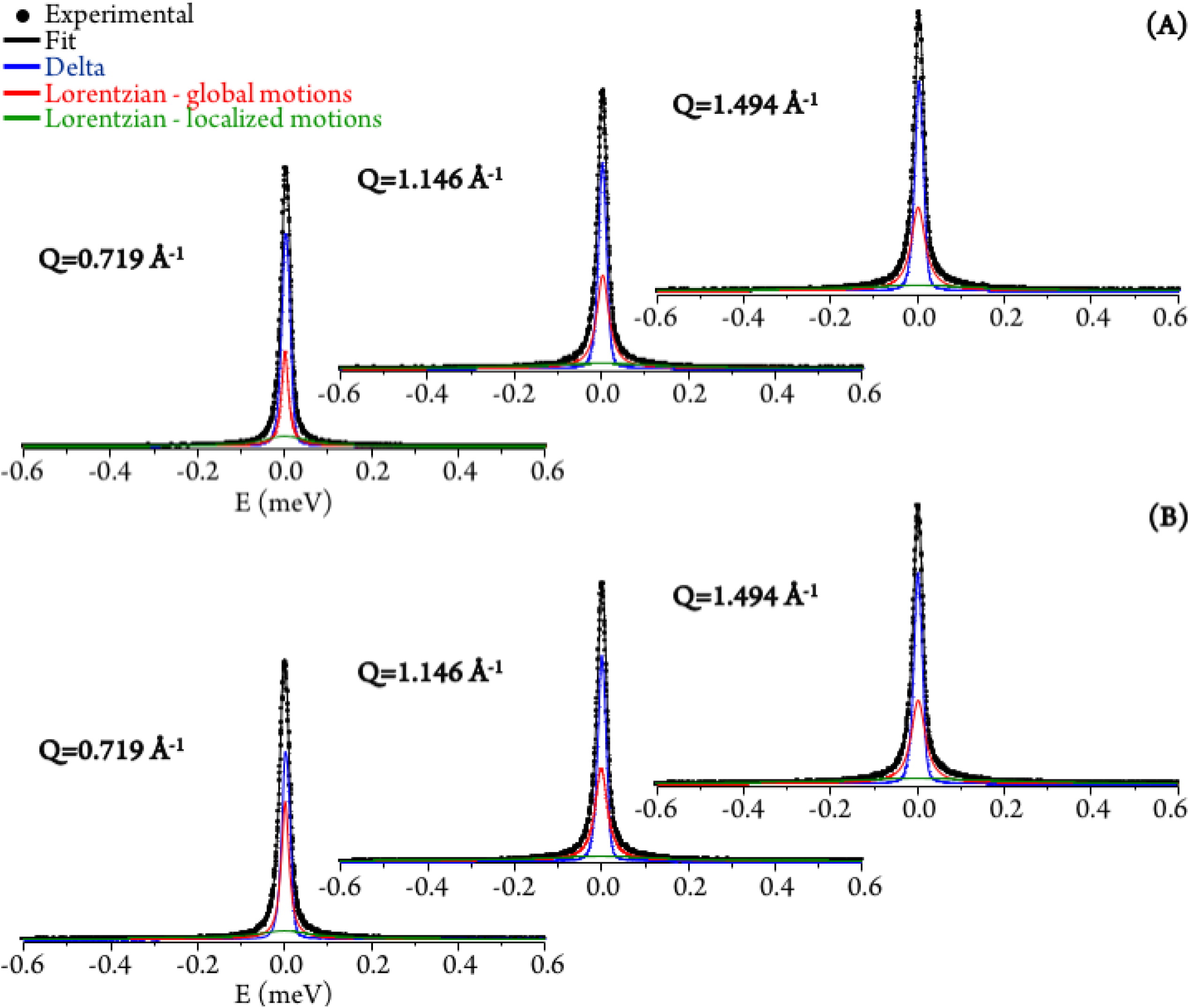
QENS spectra (298 K) for untreated (A) and Pd_2_Spm-treated/8 μM (B) H_2_O-hydrated DNA, fitted using two Lorentzians and one Delta functions, at some typical Q values.

The Q-dependent dynamical processes were consistent with a jump-diffusion reorientation model (*via* large-amplitude cooperative jumps, see Supporting Information),(Laage, 2009, Laage & Hynes, 2006) as reflected in the plots depicting the corresponding FWHM (Γ_global_) as a function of Q^2^ for both treated and untreated H_2_O-hydrated DNA, that show an asymptotic approach to a plateau at high Q values (Figure 6(A)). This behavior agrees with the water distribution in the first hydration layer of nucleic acids, predominantly binding to their outer surface (unlike for proteins were it may be enclosed between polypeptide layers or in hydrophobic pockets) and therefore allowing the formation and breaking of H-bonds between the biopolymer and its neighboring waters. In turn, the fast motions of the nucleic acid’s amine and hydroxyl groups (not restricted by H-bonding) were found to be independent of the scattering vector (Figure 6(B)), evidencing a localized dynamical process with relaxation times given by τ=(Γ_local_)^-1^.

**Figure 6:**
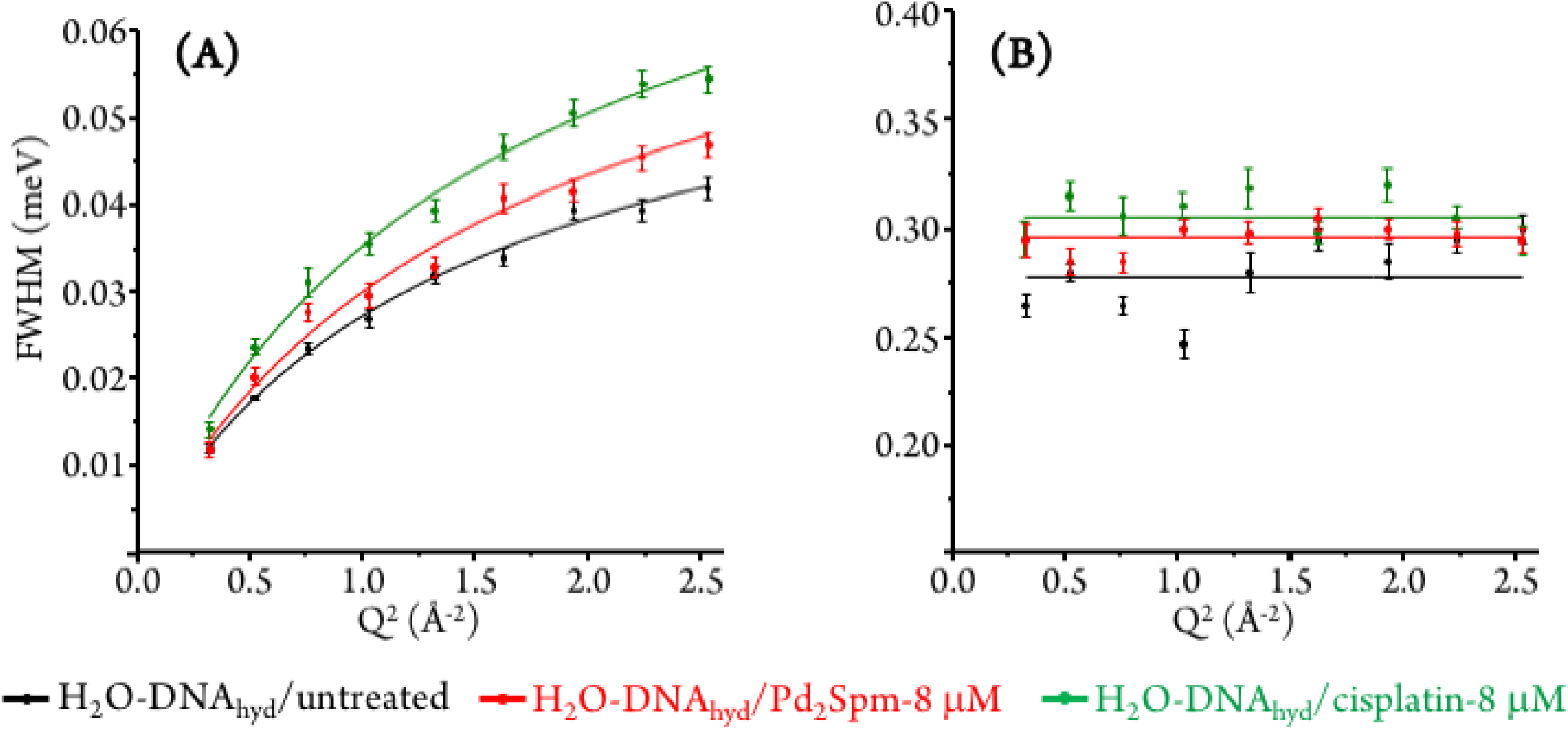
Variation of the full widths at half-maximum (FWHM) with Q^2^ for untreated, and Pd_2_Spm- and cisplatin-treated H_2_O-hydrated DNA (at 298 K): (A) Lorentzian function representing the slow translational motions (Γ_global_). (B) Lorentzian function representing the fast localised motions (Γ_local_).

Table 1 comprises the values presently obtained for the translational jump-diffusion coefficients (D_T_) and residence times between jumps (τ_T_), as well as for the correlation times of the fast localized processes, for each system studied. These results reflect the effect of the tested Pt- and Pd-agents on DNA’s dynamical behavior. For the dynamical processes associated to the hydration layer, an increase of the diffusion coefficient (7.21 to 8.95 x10^-5^ cm^2^s^-1^) coupled to a decrease of the residence time (10.1274 to 7.3974 ps) for control *versus* Pd_2_Spm- and cisplatin-exposed DNA reveal a drug-triggered enhanced mobility. The same trend was observed for the local (faster) motions, unveiled by a decrease in the corresponding correlation time (3.5927 to 3.2721 ps). A slightly more significant effect was observed for cisplatin as compared to the Pd-spermine agent, mainly regarding their effect on DNA’s hydration shell, as already revealed by the corresponding QENS profiles.

The τ_T_ value (10 ps) currently obtained for the dynamical processes taking place within DNA’s first hydration layer, in the absence of drug, agrees with previously reported data(Duboue-Dijon et al., 2016, Laage et al., 2017) and is one order of magnitude higher than the value for bulk water (1.1 ps(Bellissent-Funel, Chen et al., 1995)). This clearly evidences the restricted dynamics of water molecules in the close vicinity of DNA. This retardation relative to bulk water (up to a 6-fold slowdown at room temperature) is due to interactions of the water molecules with hydrophobic moieties and phosphate groups of the biopolymer,(Duboue-Dijon et al., 2016, Saha, Supekar et al., 2015) a small number of particularly slow water molecules having been found in DNA’s minor groove (bridging the nitrogen and oxygen atoms of complementary bases), with reorientation times between 60 and 85 ps – the so-called “spine of hydration”.(McDermott, Vanselous et al., 2017) In addition, the lower residence times currently measured for drug-incubated DNA may be partially due to a charge screening effect of the metal complexes under study (comprising partial positive charges on the Pt(II) and Pd(II) ions), that may weaken the electrostatic interactions between hydration water and negatively charged phosphates at the DNA surface, thus allowing an increased flexibility of the system. In fact, apart from the major role of hydration dynamics, electrostatic factors have also been suggested to influence the dynamical behavior of biopolymers.(Roh, 2016)

In sum, drug exposure was found to disrupt the native ordered B-conformation of DNA, prompting a higher flexibility of the nucleic acid. In the light of the results currently gathered, this is suggested to be associated to a drug impact on the biomolecule’s first hydration layer, which is prompted into a faster dynamics. This corroborates and complements the effect observed for cisplatin(Marques et al., 2017) and currently measured for Pd_2_Spm on the intra-cellular millieu, in human breast MDA-MB-231 cancer cells: cytoplasmic water was rendered more rigid by the presence of the metal complexes, while hydration water was driven into a more mobile state, as presently verified for H_2_O-DNA_hyd_ (Figure 7). Additionally, the more moderate effect on DNA dynamics revealed for Pd_2_Spm relative to cisplatin is in accordance with the influence on intracellular water that was also shown to be more significant for the mononuclear Pt-agent.

**Figure 7:**
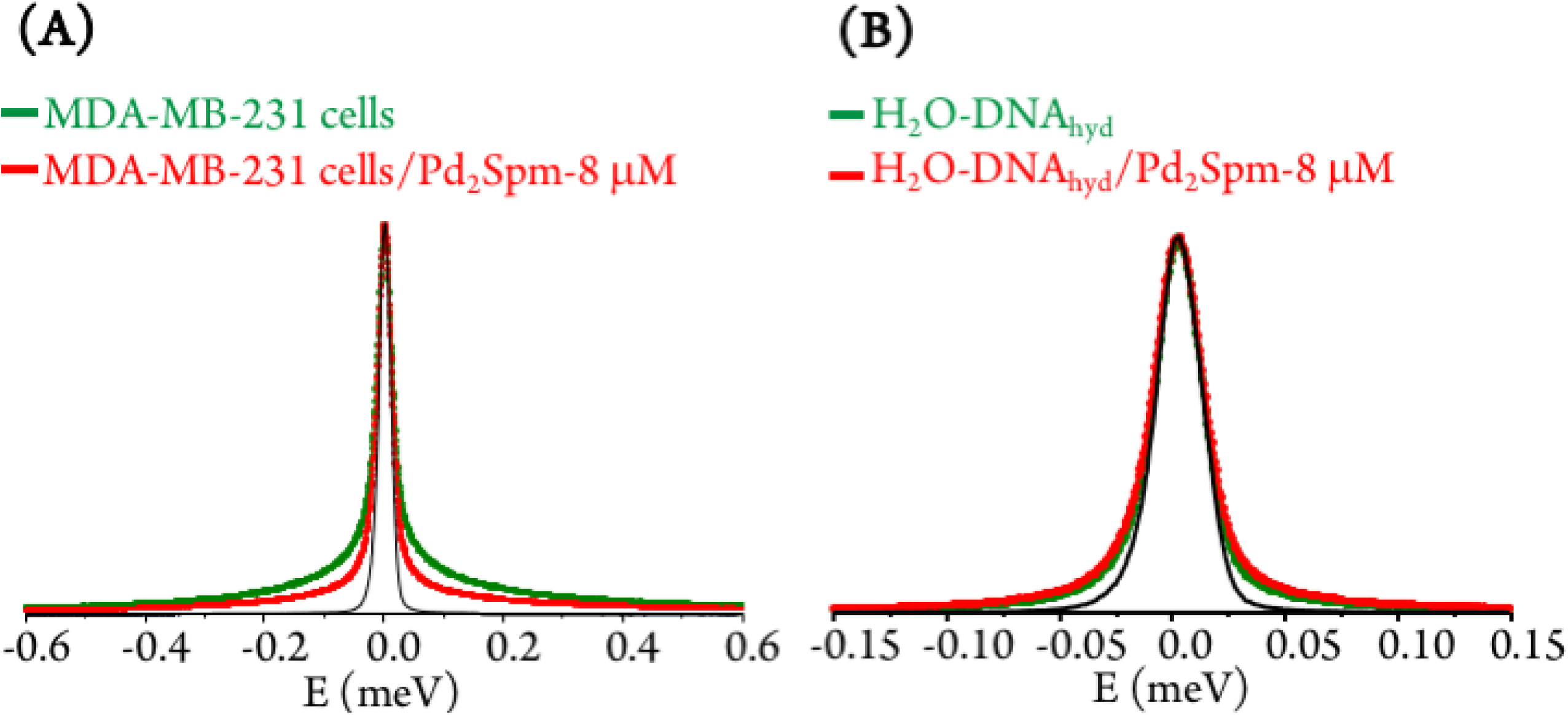
QENS data representing the drug effect on human triple negative breast cancer cells (MDA-MB-231) and on DNA: (A) QENS profiles (for all Q values) for untreated and Pd_2_Spm-treated cells. (B) QENS profiles (for all Q values) for untreated and Pd_2_Spm-exposed H_2_O-hydrated DNA. (The spectra were normalized to maximum peak intensity. The black line represents the instrument resolution).

### Synchrotron-based FTIR-ATR

SR-FTIR-ATR measurements allowed access to the spectral signatures of DNA in the absence and presence of the tested antitumor agents, thus revealing their effect on the nucleic acid’s conformation which is the basis for their cytotoxicity. The impact of the dinuclear Pt_2_Spm and Pd_2_Spm compounds was assessed, in both the far- and mid-infrared domains, the conventional mononuclear drug cisplatin having been measured for comparison purposes. Apart from the free Pt- and Pd-agents, drug-treated and untreated DNA samples were analyzed (at room temperature): for DNA extracted from MDA-MB-231 cells (DNA_pellet_) and for commercial DNA (calf thymus DNA fibers, DNA_dehyd_) taken as a reference.

This was intended as a target-oriented study on the pharmacodynamics of Pt_2_Spm and Pd_2_Spm, the vibrational fingerprint of drug-exposed DNA having been probed with a view to: (i) evaluate the drug effect on DNA – relating the induced changes in its infrared fingerprint to conformational rearrangements (*e.g.* B-DNA to A- or Z-DNA); (ii) detect each tested drug and determine its structural variations due to interaction with the nucleic acid – *via* the measured deviations in its characteristic vibrational profile, particularly in the low frequency region. To the best of the authors’ understanding, this is the first time the simultaneous detection of metal-based anticancer compounds and their pharmacological target is achieved, which constitutes an innovative way of directly monitoring a drug’s pharmacodynamics at a molecular level. Actually, although the full vibrational profile of the antitumor agents currently tackled (cisplatin, Pt_2_Spm and Pd_2_Spm) has been previously assigned by the team,(Batista de Carvalho et al., 2018, Batista de Carvalho et al., 2011, Fiuza et al., 2015, Marques et al., 2013) these compounds had never been detected within a biological matrix. This was currently made possible through the use of a high flux far-IR coherent synchrotron radiation, ensuring a high signal quality even in the low frequency region (<600 cm^-1^). Indeed, this spectral interval was particularly relevant to the study, as it allowed to simultaneously observe characteristic low energy vibrational bands from both the drug and the nucleic acid. Interpretation of the measured data was assisted by the extensive spectroscopic results formerly obtained by the authors for cisplatin, Pt_2_Spm and Pd_2_Spm,(Batista de Carvalho et al., 2018, Batista de Carvalho et al., 2011, Fiuza et al., 2015, Marques et al., 2013) as well as by reported low frequency vibrational data on DNA encompassing H-bond vibrations, base-twisting (breathing) and librational modes, known to be strongly conformation-dependent(Globus, Woolard et al., 2003, Markelz, Whitmire et al., 2002) and thus prone to be significantly affected by drug binding.

The metal complexes under study yield low frequency IR-active signals, assigned to vibrational modes involving the metal center(s): *v*(M–Cl), δ(Cl–M–Cl), δ(Cl–M–N), δ(N–M–N) and δ(N–M–Cl).(Batista de Carvalho et al., 2018, Batista de Carvalho et al., 2011, Fiuza, Amado et al., 2008, Fiuza et al., 2015, Marques et al., 2013) Some of these distinctive features were presently detected in the DNA samples extracted from drug-exposed MDA-MB-231 cells (DNA_pellet_), with clear changes relative to the bands from the isolated drugs owing to DNA coordination (Figure 8). This constitutes solid evidence of the drug’s cellular uptake and conformational rearrangement upon binding to the nucleic acid, the main observed changes being (Figures 8 and 9): (i) absence of the strong *v*(Cl–Pt–Cl) signal (*ca.* 320 cm^-1^), as expected in view of the drug activation process by intracellular chloride hydrolysis prior to interaction with the target(Marques et al., 2015); (ii) blue shift of the δ(Pt–NH) and *v*(CN/CC) signals from Pt_2_Spm and Pd_2_Spm (at *ca.* 1050 and 1070-1085 cm^-1^, respectively); (iii) shifts of the CH_2_ and NH_2_ deformation bands from the spermine ligand (at 1150-1460 and 1585 cm^-1^), more noticeable for the Pd(II) system and reflecting a conformational reorganization of the alkylamine chain upon formation of the drug-DNA adduct. Similarly, for cisplatin the NH_3_ torsion, rocking and symmetric deformation modes (at *ca.* 210, 800 and 1300-1325 cm^-1^) were found to be severely affected, which can be justified by the significantly restricted conformational freedom of the drug’s NH_3_ moieties within the DNA adducts.

**Figure 8:**
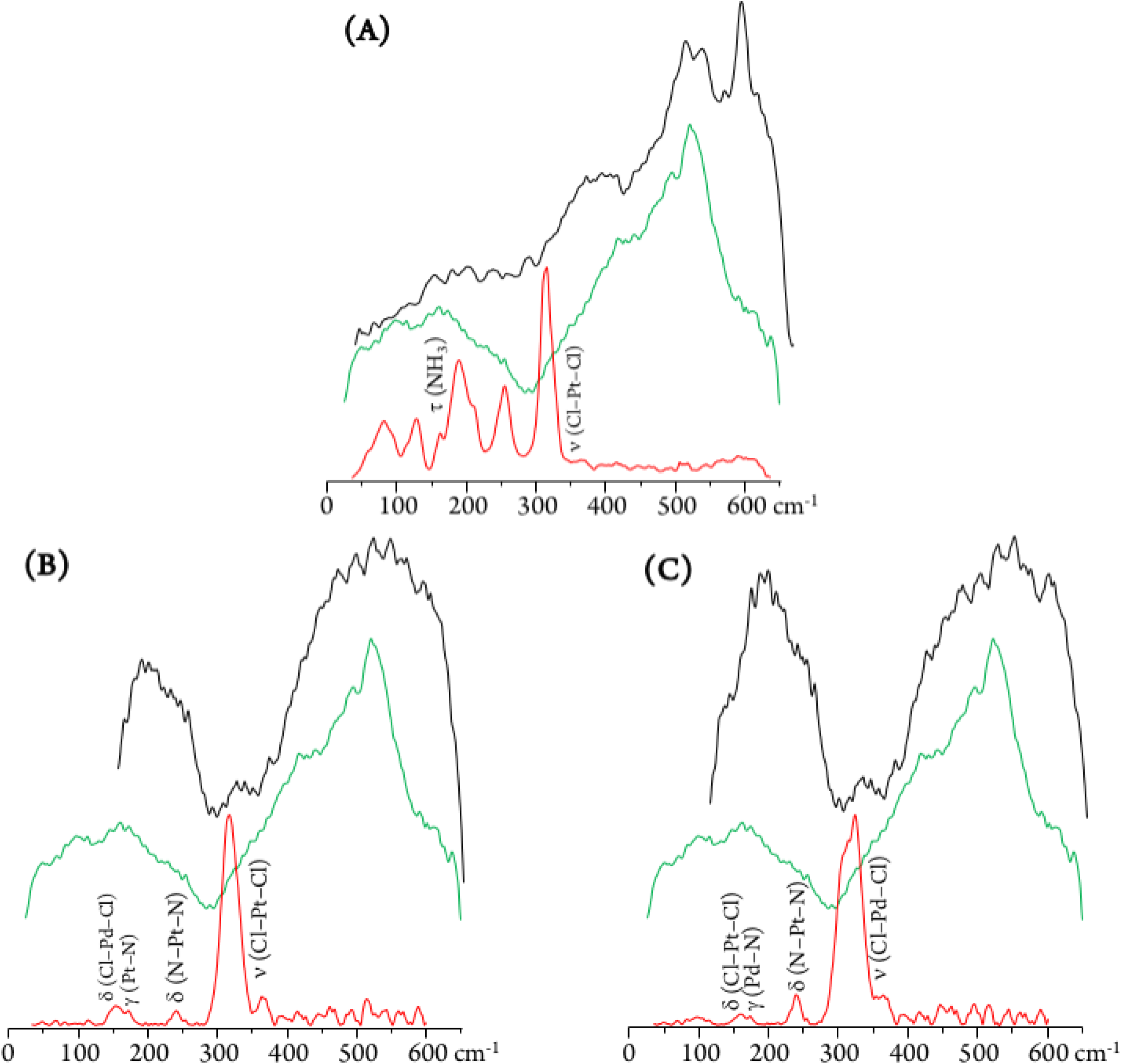
SR-farIR spectra of DNA extracted from MDA-MB-231 human breast cancer cells (DNA_pellet_): control (green), and after drug incubation (black) – cisplatin (A), Pt_2_Spm (B) and Pd_2_Spm (C). The spectra of the free drugs are also shown (red).

**Figure 9:**
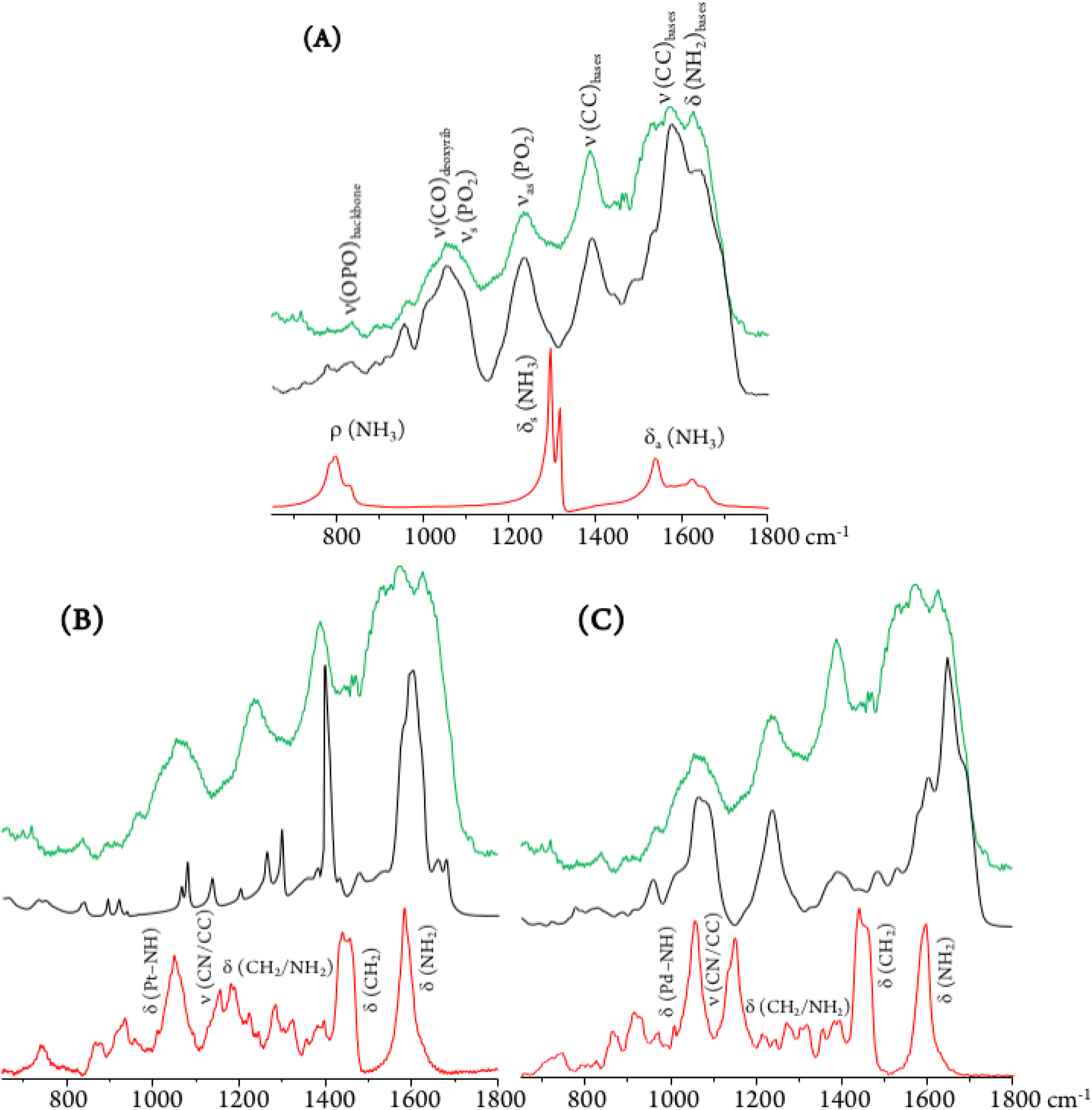
SR-midIR spectra of DNA extracted from MDA-MB-231 human breast cancer cells (DNA_pellet_): control (green), and after drug incubation (black) – cisplatin (A), Pt_2_Spm (B) and Pd_2_Spm (C). The spectra of the free drugs are also shown (red).

Regarding the main drug target, DNA, the typical phosphate stretching signals from the B-conformation – at 1092 (*v*_s_(PO_2_)) and 1237 cm^-1^ (*v*_as_(PO_2_)) – were found to undergo drug-induced deviations and intensity changes, which were much more significant upon Pt_2_Spm-treatment (Figure 9). This is indicative of a distinct type of DNA interplay for this Pt(II) agent as compared to its Pd(II) homologue, which is in agreement with previously reported evidence obtained by both FTIR and Raman microspectroscopy in drug-exposed human cancer cells.(Batista de Carvalho et al., 2016b) By comparison of the PO_2_ vibrational features for the three drugs under analysis, it is noteworthy that DNA’s phosphate groups are affected by these agents in the following order: Pt_2_Spm > Pd_2_Spm > cisplatin. In addition, the DNA samples exposed to Pt_2_Spm and Pd_2_Spm evidenced a noticeable variation of the vibrational features at 1380 cm^-1^ and 1550-1620 cm^-1^, assigned to the *v*(CC) and δ(NH_2_) modes of the double helix purine and pyrimidine bases, these bands being considerably less disturbed in the presence of cisplatin (Figure 9) which appears to reflect a distinct interaction of this compound with the bases. Actually, while cisplatin can only establish two covalent bonds with DNA’s bases (usually within the same helix), the dinucleat Pt- and Pd-spermine agents are capable of binding to four different sites simultaneously, and may yield interstrand adducts (apart from intrastrand) which cause a more severe damage.

Spectral changes were also detected in the low frequency bands ascribed to collective vibrations within double stranded DNA (phonon modes), such as helical motions, base twisting and DNA breathing (Figure 8). The latter, in particular, are associated to intrinsic fluctuational processes involving intermolecular hydrogen bonds (*e.g.* breaking of base-pair interactions creating transient DNA “bubbles”), that are highly conformation-dependent.(Gonzalez-Jimenez, Ramakrishnan et al., 2016, H., P. et al., 2013) Clear variations were measured in the spectral range 400 to 600 cm^-1^ (Figure 8), also comprising ring and carbonyl deformation modes (both in-plane and out-of-plane) from DNA bases. These are prone to be strongly affected by DNA metalation upon drug exposure, which occurs at the most nucleophilic nitrogens (N7) of the purine and pyrimidine bases. Actually, distinct spectral patterns are observed in this region for cisplatin as compared to the dinuclear agents, with a well-defined and intense band being detected at 594 cm^-1^ for cisplatin-DNA. Although this feature was also seen for the other drug-DNA pellets, it displayed a quite lower intensity suggesting a different interplay of the polynuclear agents with the double helix as compared to mononuclear cisplatin. In fact, for the spermine chelates a pre-aggregation process is known to precede the metal-target covalent binding due to a hydrophobic attraction between the drug’s polyamine ligand and DNA’s apolar backbone. This type of aggregation mechanism is absent for cisplatin. As to the spectral profile around 200 cm^-1^, it is markedly different for the cisplatin- *vs* the Pt_2_/Pd_2_Spm-containing samples. The DNA ring puckering modes assigned to this low frequency interval are overruled by characteristic water-breathing vibrations, mainly in the samples exposed to the spermine complexes (Figure 8). The region below *ca.* 100 cm^-1^, in turn, encompasses skeletal modes from the nucleic acid’s solid lattice.

The spectroscopic data currently gathered is a clear evidence of the strong interaction between the Pt(II) and Pd(II) agents presently probed and DNA – *via* metal coordination to its nitrogen bases upon a hydrophobic-driven pre-aggregation between the drug and DNA in the case of the polyamine chelates. This interplay is reflected in the noticeable narrowing of the vibrational bands measured for the drug-treated DNA *vs* the control (Figure 9), which reflects a significantly lower flexibility of the former. Furthermore, since binding of these metal-based agents to DNA triggers a severe damage of the nucleic acid’s native helical B-conformation, mainly through disruption of the double helix H-bonded base-pairs (DNA unzipping, at temperatures below the DNA melting value), the low energy intrahelical vibrational modes ascribed to DNA breathing (closed–open base-pair fluctuations) were visibly affected. These low frequency DNA modes (<300 cm^-1^), that are critical to biological function, display a marked cooperative nature and are particularly sensitive to structural reorientations, thus allowing to identify different conformational states of the biomolecule (*e.g.* B- *vs* A- or Z-DNA). Hence, the data currently reported offers unique information on the drug’s impact on DNA and helps to establish a connection between the observed low frequency vibrational fingerprint of the nucleic acid in the presence of each compound and DNA’s conformational changes that underlie cytotoxicity.

In addition, it should be emphasized that the FTIR pattern obtained for the commercial DNA_dehyd_ sample differs from that of the DNA pellets, as expected attending to the change in the hydration degree of the biomolecule.

These results, including complementary data regarding the lower frequency spectral range not previously probed, are in good accordance with the information formerly obtained (by EXAFS, FTIR, Raman and INS) for DNA’s purine bases upon cisplatin binding.(Batista de Carvalho et al., 2016b, Marques et al., 2015) In addition, valuable information was gathered on a different interplay with the nucleic acid regarding the dinuclear agents Pt_2_Spm and Pd_2_Spm as compared to the mononuclear (clinically used) drug cisplatin.

### EXAFS

EXAFS/XANES analysis provided a molecular picture of drug-DNA interplay for Pt_2_Spm and Pd_2_Spm, specifically regarding their interaction with DNA purine bases recognized to be the main binding sites for cisplatin-like agents. The major goal of the proposed study was attained: to unequivocally determine the local environment of the absorbing Pt(II) and Pd(II) centers in the drugs’ adducts with adenine and guanine (1:4), with a view to determine the precise first coordination sphere content in these entities. Additionally, interaction with glutathione was assessed, since this ubiquitous sulfur-containing tripeptide can compete for the drug relative to its major pharmacological target (DNA), thus being implicated in acquired resistance to metal-based chemotherapeutic drugs.(Chen & Kuo, 2010) The current experiments were performed for both the solid samples and the aqueous solutions, the results thus obtained showing no significant differences in accordance with previous EXAFS measurements for the homologous cisplatin-adducts (Marques et al., 2015). Regarding the effect of temperature, it was verified that the data at 90 K agreed well with that measured at room temperature, corroborating the stability of these spermine complexes under physiological conditions, a major requirement for their *in vivo* application as anticancer agents.

Interpretation of the data was based on spectra measured for solid PtA_4_, PdA_4_, PtGS_4_ and PdGS_4_, taken as references for the metal center with distinct first coordination spheres, respectively (4N) and (4S). EXAFS fit of these standards allowed us to optimize the bond length values and obtain the amplitude (S_0_^2^), energy shift (E_0_) and Debye-Waller (σ^2^) factors that were then used for fitting the metal adducts (displaying coordination environments with either equal or different ligands).

The parameters of the nearest coordination shell around Pd and Pt were obtained, namely the number and type of neighbor atoms, and their distance from the selected metal (Table 2). It was clearly observed that stoichiometric amounts of either adenine or guanine led to a complete coordination to the metal centers in both the Pt and Pd adducts – MA4 and MG4, displaying a (4N) environment *per* metal as opposed to (2N+2Cl) in the free complexes (Figure 10 (A) and (B)). When comparing both purine bases as binding sites for the Pt_2_Spm and Pd_2_Spm agents, the symmetry was found to be significantly higher for the Pt(II) adducts (reflected by the stronger inten sity of the EXAFS signals). When comparing the Pt(II) and Pd(II) adducts with DNA’s purine bases, the latter displayed slightly shorter Pd–N values in both the adenine and guanine systems (respectively 2.027±0.007 *vs* 2.039±0.005 and 2.035±0.006 *vs* 2.047±0.005 Å, Tab. 2). In addition, it should be noted that the Pd–N bond lengths within PdA4 and PdG4 were also shortened relative to the Pt(NH_3_)_4_ reference sample previously measured (Marques et al., 2015), (2.039±0.004 Å).

**Figure 10:**
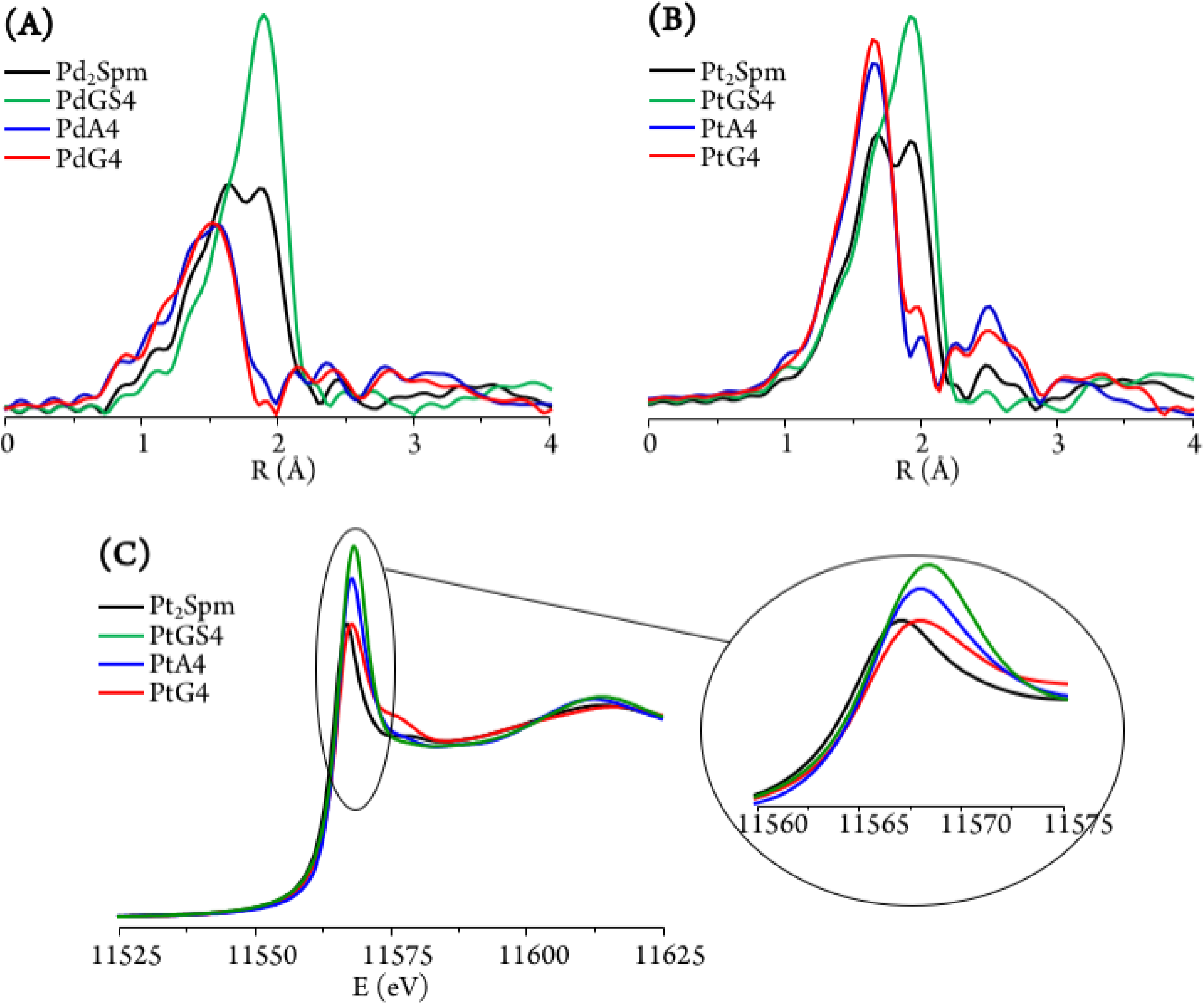
Fourier transform of EXAFS χ(k) data (at room temperature) and best fits (A and B), and XANES profiles (C) for Pt_2_Spm, Pd_2_Spm and their (1:4) adducts with adenine (MA4), guanine (MG4) and glutathione (MAS4 and MGS4) (M=Pt(II) or Pd(II)).

For the drug-glutathione (1:4) adducts, a (4S) first coordination shell was detected for each of the metal centers (Figure 10(A) and (B), Table 2), as proof of an efficient drug binding to glutathione molecules (monodentate, solely *via* GSH’s S atoms). Both for the Pt- and Pd-glutathione adducts, the calculated M–S bond lengths compare well with that obtained for the Pt(GSH)_4_ reference (Marques et al., 2015) (2.308±0.006 and 2.334±0.004, respectively, *vs* 2.313±0.006 Å). PdGS4, however, displays a somewhat longer metal-to-sulfur distance, revealing a slightly weaker glutathione coordination.

The slight shift of the edge (to higher values) in the XANES profiles observed for the distinct systems under study evidenced a variation in the degree of electron donation from the ligand atoms to the metal center (Table 3). While this charge transfer process was similar for both Pd(II)-purine adducts differing only for PdGS4, for the Pt(II) systems this ligand-to-metal electron delocalization was found to increase from the free complex ((2N+2Cl) environment) to the PtA4/PtG4 ((4N)) and PtGS4 ((4S)) adducts, as expected attending to the increasingly softer nature of the surrounding ligands. This fact has relevant implications regarding the glutathione-induced resistance associated to this type of metal-agents. Furthermore, the marked intensity increase detected for PtA4 and PtGS4 (Figure 10(C)) reflects a significantly lower symmetry of these species as compared to PtG4 (and to the free drug), since this pre-edge intensity is strongly dependent on the size and geometry of the molecular cage surrounding the metal absorber. For the Pd(II) systems, in turn, this symmetry loss was hardly noticeable, which may suggest a more labile coordination of this cation relative to Pt(II), the latter displaying a clear preference for guanine – yielding PtG4 stable adducts with a defined and quite rigid geometry.

Apart from the purine and glutathione adducts of Pt_2_Spm and Pd_2_Spm, the drug’s preference for glutathione *versus* DNA binding sites (purine’s N7 atoms) was assessed. For the PdA4 and PdG4 systems, the presence of glutathione (1:4) was found to lead to a N by S substitution, a (4N) metal coordination sphere giving rise to *ca.* (3N+1S) and (4S) patterns for the metal centers within the adenine and guanine adducts, respectively (Figure 11(A), Table 2). This clearly evidences a lower stability of the Pd_2_Spm-guanine adducts, for which there was a total substitution of the nitrogen ligand atoms by sulfur, leading even to a disruption of the Pd_2_Spm complex (Pd(II) being detached from the spermine NH_2_ ligand moieties, at both metal centers). Interestingly enough, this is contrary to what had been previously found for the mononuclear cisplatin-A4 and cisplatin-G4 systems, the latter having shown to be stabilised(Marques et al., 2015) (as expected, through an intramolecular NH _drug_ **^…^**O=C _guanine_ hydrogen close contact). A possible justification for the behavior presently found for the dinuclear spermine agents may be the less favorable H-bond between their amine groups and the guanine’s carbonyl as compared to cisplatin, coupled to a higher steric hindrance between the Pd_2_Spm-base adducts and glutathione on account of the bulkier polyamine ligand and the presence of two metal centers. The Pt_2_Spm-base adducts, in turn, revealed a quite distinct behavior relative to their Pd homologues in the presence of glutathione, both the adenine and guanine systems displaying a similar degree of S-substitution yielding an approximate (1N+3S) shell (Figure 11(B)). Overall, PdA4 was shown to be the most stable adduct relative to glutathione competition, only one nitrogen ligand having been substituted by sulfur. These results were confirmed by the XANES linear combination fits, comprised in Table 3. Upon GSH coordination to the drug adducts, the M–S bond lengths were found to be identical for all systems under study (2.31 to 2.33 Å, Table 2). Moreover, S-binding did not affect the metal-nitrogen coordination within the adducts, the M–N values being equal for PtA4+GSH, PtG4+GSH and PdA4+GSH (2.03 Å, Table 2).

**Figure 11:**
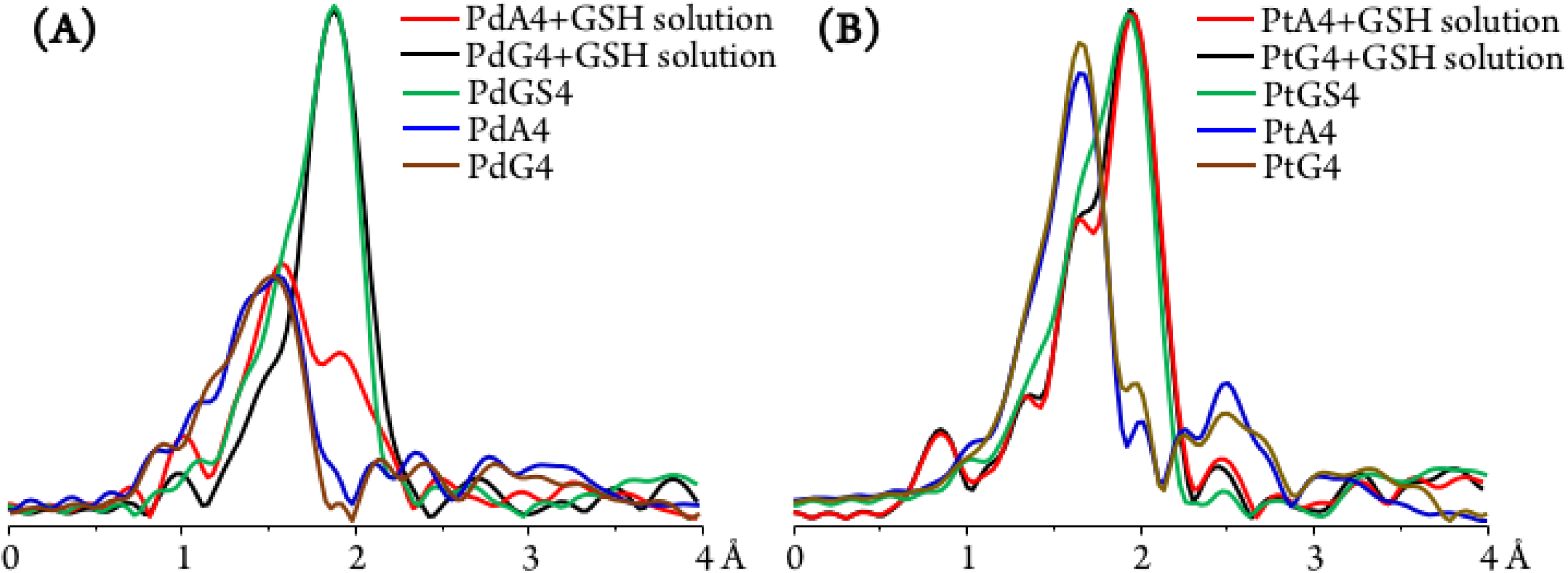
Fourier transform of EXAFS χ(k) data (at room temperature) and best fits for the systems obtained when adding glutathione (1:4) to MA4 and MG4 solutions (M=Pt(II) or Pd(II)): (A) Pd_2_Spm; (B) Pt_2_Spm. (The data measured for each of the pure adducts is included for comparison).

Glutathione competition for the drugs, well justified by the higher affinity of Pt(II) and Pd(II) for GSH’s sulfur over adenine’s and guanine’s nitrogens, has been previously found for cisplatin(Marques et al., 2015) and constitutes experimental proof of glutathione-mediated drug inactivation (underlying acquired resistance) also for the dinuclear agents presently investigated.

## CONCLUSIONS

The dinuclear complexes Pt_2_Spm and Pd_2_Spm have been studied by the authors in the last few years and have been shown to act as promising anticancer agents towards human metastatic breast cancer(Batista de Carvalho et al., 2016a, Batista de Carvalho et al., 2016b, Fiuza et al., 2011, Marques et al., 2002, Silva et al., 2013) and osteosarcoma(Lamego et al., 2017). A multidisciplinary study was presently carried out to attain complementary structural and dynamical information on their interplay with DNA (their main pharmacological target), using QENS measurements, SR-FTIR and SR-EXAFS/XANES.

The QENS experiments currently performed provided accurate and unique data on the impact of metal-based anticancer drugs on DNA *via* the biopolymer’s first hydration layer, which is known to be closely coupled to DNA function. Indeed, any drug-triggered perturbation on the nucleic acid’s hydration shell is expected to influence the biomolecule’s dynamical profile and consequently its biological function. Measurements for drug-treated and untreated H_2_O-hydrated DNA, at room temperature, revealed a clear effect of both cisplatin and Pd_2_Spm on the nucleic acid’s hydration shell dynamics, reflected in an increased flexibility for the drug-incubated samples. Two dynamical processes were discriminated (in the ps timescale), ascribed to: (i) the biopolymer’s hydration water; (ii) fast localized rotations of specific groups within DNA – CH_3_, as well as NH_2_ not restricted by hydrogen-bonds. The intrinsic stiffness of native DNA appears to be disrupted by the drug, underlining the influence of the water molecules from the first hydration sheath. It should be emphasized that this effect was only observed (in the ps timescale) for H_2_O-hydrated DNA and not for the D_2_O-hydrated nucleic acid, which also evidenced distinct dynamical transition temperatures. Apart from the hydration-induced flexibility increase, an electrostatic-mediated dynamic enhancement was unveiled for drug-exposed H_2_O-hydrated DNA. These conclusions are in good agreement with the previously reported drug impact on the intracellular millieu – cytoplasmic water and biomolecules’ hydration layers:(Marques et al., 2017) while the former showed a restrained dynamics upon drug exposure, the hydration shells were prompted into a more mobile state, as presently corroborated for DNA. These combined results support the pronounced effect of metal-based chemotherapeutic agents on water dynamics, and should help to gain a more thorough understanding of the drug-induced cytotoxicity *via* the water molecules both in the intracellular medium and in the close vicinity of biopolymers. In addition, an interesting onset of anharmonicity was detected for dehydrated DNA (in the ps timescale) in contrast with previous studies on dehydrated biopolymers (*e.g.* proteins, RNA and DNA), and it was tentatively assigned to rotational motions of methyl groups (1 *per* 4 bases within the double helix) and, to a smaller extent, of H-bond free NH_2_ moieties.

Infrared spectroscopy with synchrotron radiation was confirmed as a powerful non-invasive molecular probe of biosamples, allowing to interrogate and obtain an accurate insight on the biochemical impact of chemotherapy drugs at the cellular level. Following previous studies on the effect of the presently tested compounds in cancer cells,(Batista de Carvalho et al., 2016b) their impact on DNA was currently probed, for DNA samples extracted from treated and untreated human MDA-MB-231 cells. The twofold purpose of this study was accomplished, since the simultaneous detection of the drug and its pharmacological target was achieved, specific spectral biomarkers having been identified reflecting changes in both the metal agent and DNA, prompted by drug-target interaction. Furthermore, the vibrational modes comprised in the far-infrared region, which are a spectral signature of the nucleic acid conformational state, allowed to unveil drug-elicited perturbations in this biomolecule recognized to be the basis of cytotoxicity: an altered pattern was detected for the bands ascribed to DNA breathing, as a consequence of a drug-triggered disruption of the double helix H-bonded base-pairs. To the best of the authors’ knowledge, this is the first experimental observation of the DNA breathing process in the presence of a chemotherapeutic agent. Additionally, analysis of the variations occurring in the characteristic bands of the complexes provided evidence of structural variations upon accumulation at the biological target. Moreover, sampling DNA extracted from drug-exposed human cancer cells enabled to qualitatively probe the drug’s cellular uptake and its activation process (through intracellular chloride hydrolysis) prior to DNA interaction, apart from its impact on the nucleic acid upon binding to the purine bases. These results could only be successfully achieved through FTIR spectroscopy with a coherent synchrotron radiation that guarantees both broadband spectral coverage and a high signal quality. The SR-FTIR results presently gathered complement previous SR-based infrared data obtained at the MIRIAM beamline for the same compounds and cell line (at equal concentrations and incubation time):(Batista de Carvalho et al., 2016b) comparing these results on whole cells with those measured for DNA isolated from cells pre-incubated with the chemotherapeutic agents is of key biological significance for discriminating drug effects regarding other potential cellular targets (*e.g.* DNA *vs* proteins, membrane lipids or intracellular water).

The successful EXAFS/XANES results gathered for the adenine, guanine and glutathione adducts with Pt_2_Spm and Pd_2_Spm allowed an accurate assessment of their pharmacodynamics (interaction with DNA) and pharmacokinetics: the local environment of the metal centers was obtained, for each system, and GSH-metal binding (solely *via* glutathione’s sulfur atom) was clearly evidenced thus justifying the glutathione-mediated drug resistance mechanisms occurring *in vivo*. A slightly weaker glutathione coordination was unveiled for Pd_2_Spm as compared to Pt_2_Spm, which may be relevant with a view to overcome *in vivo* GSH-mediated drug resistance. Additionally, EXAFS measurements for the adducts in the presence of glutathione revealed a lower stability of the Pd_2_Spm-guanine adducts relative to their adenine homologues (contrary to what had been previously found for cisplatin(Marques et al., 2015)), while their Pt counterparts showed to have an identical stability to the mononuclear clinical drug. Coupled to the data previously reported for cisplatin,(Marques et al., 2015) these results provided accurate information on the chemical composition, structure and relative stability of the adducts formed between the Pd_2_Spm and Pt_2_Spm complexes and DNA purine bases, leading to a better understanding of the mode of action of this type of polynuclear metal agents at the molecular level.

The present multidisciplinary study on the impact of cisplatin and cisplatin-like dinuclear agents on DNA’s dynamical and structural profiles constitutes an innovative way of tackling a drug’s mode of action. A comprehensive and reliable set of data was gathered on the molecular basis of their cytotoxicity towards the very low prognosis human metastatic breast cancer, allowing to better clarify the unconventional interaction of Pt_2_Spm and Pd_2_Spm with their main target. Combined with biological assays for evaluation of antitumor activity, the current results are expected to provide valuable clues for the rational design of metal-based compounds with improved therapeutic properties, acting through a multistep process that leads to function loss of vital biomolecules and ultimately to cell death: (i) direct binding to DNA, causing disruption of its native conformation and prompting biofunctional disability; (ii) continuing effect on the nucleic acid’s hydration layer, which is prompted into a faster dynamics, triggering changes in the biopolymer with consequences at the functional level; (iii) impact on intracellular water (cytosol), with an expected global effect on essential cellular components thus hindering normal cellular function.

## MATERIALS AND METHODS

The list of chemicals, the experimental details regarding the synthesis and characterization of the Pt_2_Spm and Pd_2_Spm complexes, the preparation of drug solutions, and the cell culture protocol are extensively described in the Supporting Information, as well as the pre-processing and analysis procedures of the QENS, FTIR-ATR and EXAFS/XANES data.

### Synthesis and Characterization of the Drug–Purine and Drug–Glutathione Adducts

*Drug-Purine Adducts:* The (1:4) drug-adenine and drug-guanine adducts (MA4 and MG4, M=Pt(II) or Pd(II)) were synthesized following a synthetic route previously optimized by the authors(Lippert, Raudaschl et al., 1984, Schoellhorn, Raudaschl-Sieber et al., 1985) (see details in the Supporting Information).

*Drug-Glutathione Adducts:* In order to prepare (1:4) drug-GSH adducts, 4 molar equivalents of glutathione were added to a solution of each of the drugs – Pt_2_Spm-0.45 mM or Pd_2_Spm-0.25 mM – and stirring was maintained in the dark, at room temperature, for 24 h. The initial colorless reaction mixture became orange over time (a characteristic color of complexes containing Pt-S covalent bonds). The resulting drug-GS4 solutions (MGS4, M=Pt(II) or Pd(II)) were then concentrated by rotary evaporation under vacuum and lyophilized to obtain an orange powder.

*Titration of Drug-Purine Adducts with Glutathione, Adenine or Guanine:* The Mbase4 (M=Pt(II) or Pd(II), base=A or G) adducts were titrated with GSH. A glutathione solution was added, in a (1:4) molar ratio, to an aqueous solution of 10 mg of each Mbase4 adduct. The reaction mixtures were kept in the dark with stirring, at 30 °C for 24 hours, after which they were concentrated under vacuum, at 30 °C until precipitation occurred and then vacuum filtered through a 0.22 μm filter to remove precipitated adenine or guanine. Finally, they were concentrated under vacuum (at 30 °C) and lyophilized to yield the solid products (hereafter denominated MA4+GSH and MG4+GSH).

### Preparation of Cell Pellets for QENS measurements

Cell pellets (100 mg/1 cm^3^, *ca.* 5×10^8^ cells *per* sample), were prepared by cell harvesting (through trypsinization) followed by repeated (2×) PBS washing and centrifugation (at 195× g, for 15 min). PBS was used as an isotonic medium in order to avoid water exchange from the inside to the outside of the cell (leading to cell shrinkage). The drugs (cisplatin, Pt_2_Spm and Pd_2_Spm at 4 and 8 μM) were added to the cells during their logarithmic phase of growth, and left to incubate for 48 h. In order to completely remove the extracellular water component (less than 5%), the cell pellets were washed with deuterated PBS by resuspension (1×) followed by centrifugation (at 195× g) for 5 min, which was then repeated for 15 min (after removal of the first supernatant).

### DNA Extraction and Preparation of Drug-DNA Samples

DNA was extracted from MDA-MB-231 cells (henceforth denominated DNA_pellet_), both untreated and treated with either cisplatin or Pd_2_Spm (see details in the Supporting Information). Fibrous commercial human DNA (from calf-thymus) was also analyzed – samples without and with drug were prepared, the latter by solubilizing 100 mg of DNA fibers in 50 mL of either cisplatin, Pd_2_Spm or Pt_2_Spm at 8 μM, with stirring (at 4° C) for *ca.* 24 h. Aqueous drug solutions were used (instead of saline solutions) in order to ensure a prompt hydrolysis of the chlorides, that is essential for drug activation prior to DNA binding. 5 mL of 3 M-sodium acetate and 150 mL of ethanol (≥99.8%) were then added, followed by a 2 h incubation (at -20°C). The solutions were centrifuged at 4075×g for 20 min (4° C), the pellets were washed twice with ethanol (70%) and centrifuged again (under the same conditions). After discarding the supernatant, the obtained DNA samples were air dried completely in a desiccator – hereafter denominated DNA_dehyd_. Apart from this dehydrated DNA, H_2_O- and D_2_O-hydrated DNA were prepared at a r.h. >80% (>0.80 g H_2_O/D_2_O/g DNA), in order to ensure the stability of the native B conformation – yielding the samples henceforth denoted as DNA_hyd_ (either H_2_O-DNA_hyd_ or D_2_O-DNA_hyd_). This was achieved by placing DNA_dehyd_ in a desiccator (closed environment) with a saturated KCl solution (in either H_2_O or D_2_O) until attaining a stable weight (corresponding to a r.h. of 84.34-83.62%, at 25 °C).(Greenspan, 1977)

### QENS Measurements

QENS data were acquired at the ISIS Pulsed Neutron and Muon Source of the Rutherford Appleton Laboratory (United Kingdom), on the OSIRIS spectrometer(Telling & Andersen, 2005) (see details in the Supporting Information). H_2_O-DNA_hyd_ and D_2_O-DNA_hyd_ samples were measured, as well as dehydrated DNA (DNA_dehyd_), both untreated and drug-exposed – to cisplatin or Pd_2_Spm at 8 μM (for a 48 h incubation time). The samples were mounted in indium-sealed 0.1 mm-thick (3×5 cm) flat Al cans (the beamsize at the sample being 2.2×4.4 cm), and were oriented at -30° with respect to the incident beam. A vanadium sample (purely incoherent elastic scatterer) was also measured, to define the instrument resolution and correct for detector efficiency. Experiments were carried out at 150 and 298 K.

### Synchrotron-based FTIR-ATR Measurements

FTIR-ATR spectra were recorded at the MIRIAM beamline B22 of DLS,(Cinque, Frogley et al., 2011) in a Bruker Vertex 80v Fourier Transform IR interferometer, in both the far-IR (FIR) and mid-IR (MIR) ranges (see details in the Supporting Information). Three sets of experiments were carried out (at room temperature): 1) pure drugs (solid powders); 2) DNA control samples, in the absence of drug – both commercial (DNA_dehyd_ fibers) and extracted from untreated MDA-MB-231 cells (DNA_pellet_); 3) drug-containing DNA samples, extracted from drug-treated MDA-MB-231 cells (drug-DNA_pellet_).

### EXAFS Experiments

X-ray absorption experiments (simultaneous EXAFS and XANES) were performed (in both the solid state and in solution) at the B18 beamline from DLS(Dent, Cibin et al., 2009) (see details in the Supporting Information). Apart from the free Pd_2_Spm and Pt_2_Spm complexes, their (1:4) adducts with adenine (A) and guanine (G) were measured (MA4 and MG4), as well as the (1:4) adducts with reduced glutathione (MAS4 and MGS4) and the adducts titrated with GSH (MA4+GSH and MG4+GSH) (M=Pt(II) or Pd(II)).

## ACKNOWLEDGMENTS

The authors thank financial support from the Portuguese Foundation for Science and Technology – UID/MULTI/00070/2013 and PTDC/QEQ-MED/1890/2014 (within Project 3599-PPCDT – jointly financed by the European Community Fund FEDER). The STFC Rutherford Appleton Laboratory is thanked for access to the Research Complex at Harwell (cell culture labs) and to the neutron beam facilities. The neutron work was supported by the European Commission under the 7^th^ Framework Programme through the Key Action: Strengthening the European Research Area, Research Infrastructures (contract n°: CP-CSA_INFRA-2008-1.1.1 Number 226507-NMI3). Diamond Light Source (UK) is acknowledged for access to the B22/MIRIAM (SM14895) and B18/Core EXAFS (SP16058) beamlines. A. Fitzpatrick is thanked for technical support at B22.

## AUTHOR CONTRIBUTIONS

ALMBC, LAEBC and MPMM conceived the study and designed the workplan. Experimental work: ALMBC, APM, VGS, AD, JD, MF, DG, LAEBC and MPMM. Data analysis: ALMBC, VGS, PG, GC, DG and MPMM. Intellectual input: VGC, PG, GC, DG, LAEBC, MPMM. Manuscript writing: ALMBC and MPM

